# Monitoring Unfolding of Titin I27 single- and bi-Domain with High-Pressure NMR Spectroscopy

**DOI:** 10.1101/292359

**Authors:** I. Herrada, P. Barthe, M. Van Heusden, K DeGuillen, L Mammri, S. Delbecq, F. Rico, C. Roumestand

## Abstract

A complete description of the pathways and mechanisms of protein folding requires a detailed structural and energetic characterization of the folding energy landscape. Simulations, when corroborated by experimental data yielding global information on the folding process, can provide this level of insight. Molecular Dynamics (MD) has been associated often to force spectroscopy experiments to decipher the unfolding mechanism of titin Ig-like single- or multi-domain, the giant multi-modular protein from sarcomere, yielding information on the sequential events during titin unfolding under stretching. Here, we used high-pressure NMR to monitor the unfolding of titin I27 Ig-like single-domain and tandem. Since this method brings residue-specific information on the folding process, it can provide quasi-atomic details on this process, without the help of MD simulations. Globally, the results of our high-pressure analysis are in agreement with previous results obtained by the association of experimental measurements and MD simulation and/or protein engineering, although the intermediate folding state caused by the early detachment of the AB ß-sheet, often reported in previous works based on MD or force spectroscopy, cannot be detected. On the other hand, the A’G parallel ß-sheet of the ß-sandwich has been confirmed as the Achilles heel of the 3D scaffold: its disruption yields complete unfolding, with very similar characteristics (free energy, unfolding volume, kinetics constant rates) for the two constructs.

## INTRODUCTION

Titin is an intra-sarcomeric protein of striated muscle that plays complex roles in myofibril mechanics and mechano-sensing (1–2). It is a giant protein (> 3 MDa) that spans *in situ* one-half of the sarcomere (> 1 µm), the repeating contractile unit of muscle. The N-terminus is at the boundary of sarcomere, the Z-disc, while the C-terminus is in the M-line at the center of the sarcomer (3). More than one-half of the molecule (the A-band) is an integral part of the myosin filament. This region is formed mainly by immunoglobulin-like (Ig) and fibronectin type III (fn3) domains. The I-band of titin forms an elastic connection between the end of the thick filament and the Z-disc. This part of titin does not contain Fn3 domains but Ig domains. In the I-band, most of the Ig domains are arranged in tandem – that is without linkers between them – in two segments, “proximal” and “distal” to the Z-line, which flank a PEVK-rich motif (4). Prior to establishing the key role of the PEVK region of titin I-band in passive elasticity of striated muscle, Ig stretch-unfolding was thought to drive the mechanical response of titin. This is now a controversial concept and Ig unfolding is no longer regarded as a primary mechanism of titin elasticity in vivo (5,6), but rather as a safety mechanism to limit high forces in the sarcomere (7–8). On the other hand, using Fluorescence microscopy combined with titin specific quantum dot labels, Rivas-Pardo et al. (9) have recently shown that passive stretching of muscle tissue stores elastic energy by unfolding Ig domains, and that refolding of these domains supplies an important component of the force delivered by a contracting muscle.

All these findings have motivated extensive studies on Ig unfolding: their thermal, chemical and mechanical stability have been investigated (10–12). These measurements often concerned Ig single-modules, and more specifically I27 (also called I91), the 27^th^ Ig-like domain in the I-band, more rarely Ig tandems or multi-modules (13–17), using different techniques as fluorescence or circular dichroism spectroscopy. Also, mechanical unfolding with atomic force microscope (AFM) has been used for I27 domain repeats up to 12 modules (14,15, 18). Even if these techniques are able to bring information on the folding landscape (is there any folding intermediate?), the transition state ensemble (is its hydration state close to that of the native state or to the unfolded state?) explored by a protein during the folding/unfolding reaction, none is able to reveal atomic details on the corresponding landscape. Likewise, while force spectroscopy experimental studies conveyed how far titin can be stretched and how much force it can bear, they cannot ascertain what specific molecular mechanisms were responsible for granting titin its mechanical resilience. Thus, these experimental techniques should be coupled to MD simulations and/or protein engineering to shed light on the missing details (15,18-20; for a review 21).

NMR spectroscopy has emerged as a particularly powerful tool to obtain high-resolution structural information about protein folding events because an abundance of site-specific probes can be studied simultaneously in a multidimensional NMR spectrum. During the last past years, we have shown that the combination of high pressure and NMR constitutes a powerful tool that can yield unprecedented details on protein unfolding landscape (22-26; for a recent review 27). Indeed, the atomic resolution offered by multidimensional NMR experiments provides an intrinsic multi-probe approach to assess the degree of protein folding cooperativity, which is otherwise difficult to characterize using techniques such as circular dichroism or fluorescence. In addition, the high reversibility of pressure unfolding/refolding experiments ensures a proper thermodynamic characterization of the process, which is often problematic to assess by heat denaturation, for instance, because of excessive aggregation. Under the influence of high hydrostatic pressure, water molecules are believed to penetrate into internal cavities of the protein core and to induce the destabilization of hydrophobic interactions [23,28]. Because solvent-excluded cavities are not uniformly distributed but rather depend on the unique structural characteristics of a protein structure, the pressure-induced unfolding originates from specific, local and unique properties of the folded state. In this sense, it is very different from unfolding by temperature or chemical denaturants, which act globally and depend on exposed surface area in the unfolded state.

In the present study, we used high-pressure NMR spectroscopy to monitor the unfolding of titin I27 as a single-module or as a tandem. This technique allowed us to reveal structural details on the folding pathway of these two constructs that are globally in agreement with previous results obtained with MD simulation coupled to protein engineering or force spectroscopy.

## MATERIALS AND METHODS

### Protein expression and purification

The constructs (subcloned in pdbccdb 3C His) allowing the expression of titin single-domain and tandem were transformed into E. coli BL21(DE3). After protein expression (3 h, 37°C, then induction with 0.2 mM IPTG overnight at 30°C), cells were suspended in lysis buffer comprising 50 mM Tris-HCl buffered at pH 7.0 and containing 300 mM NaCl and 1 mM DTT. Lysozyme at 1µg/µl was added to aid cell disruption. Cells were lysed by sonication (2 s bursts for 3 min, at 60% amplitude, with a large parallel probe, Vibra cell 72405). Cell debris and insoluble materials were removed by centrifugation (Beckman Coulter Avanti J-20 XP centrifuge equipped with a 25.50 rotor, set at 18,000 rpm, at 6°C). The supernatant was filtered through a 0.45 µm PVDF filter (Millipore) and loaded through a AKTA system onto a 1 ml HisTrap HP columm (GE Healthcare) equilibrated with lysis buffer. After imidazole gradient elution, fractions containing the protein were identified by SDS-PAGE. Protein was put in dialysis membrane (ThermoFisher Scientific) with 3C protease (0.1mg/ml) at 4°C over-night in 20 mM Tris-HCl buffered at pH 7.0, 150mM NaCl and 1 mM DTT. Cleavage was checked with SDS-PAGE and the protein was finally injected through an AKTA system into a Superdex S75 16/60 (GE Healthcare) column, equilibrated with the same buffer used for dialysis. The fractions containing the pure protein were pooled, concentrated to obtain 1 mM protein (Vivaspin 20, Sartorius Stedim Biotech) then dialyzed with 20mM Tris-HCl 10 mM DTT and 0.01mM PMSF and stored at −80°C until NMR analysis. Uniform ^15^N labeling was obtained by growing cells in minimal M9 medium containing ^15^NH_4_Cl as the sole nitrogen source (Cambridge Isotope Laboratories Inc.).

### NMR assignment en solution structure

Protein (I27 single-module or tandem) samples were dissolved in 200 µL of aqueous buffer containing 20 mM Tris-HCl pH 7.0 and 10 mM DTT (5% D2O for the lock) at a concentration of about 1 mM. Except if otherwise specified in the text, all experiments were recorded at 25°C on a Bruker AVANCE III 600 MHz equipped with a proton detection broad band inverse (BBI) 5 mm Z-gradient ^1^H-X probe head. Water suppression was achieved with the WATERGATE sequence (29). ^1^H chemical shifts were directly referenced to the methyl resonance of DSS, while ^15^N chemical shifts were referenced indirectly to the absolute frequency ratio ^15^N/^1^H = 0.101329118. All NMR experiments were processed with Gifa (30).

3D [^1^H,^15^N] NOESY-HSQC (mixing time 100 ms) and TOCSY-HSQC (isotropic mixing: 60 ms) NMR double-resonance experiments were used for sequential assignment and to extract nOe’s restraints to model the solution structure. NOE cross-peaks identified on 3D [^1^H, ^15^N] NOESY-HSQC (mixing time 150 ms) were assigned through automated NMR structure calculations with CYANA 2.1 (31). The nOe’s data set was completed by the analysis of a 2D NOESY recorded at 800 MHz (AVANCE III Bruker spectrometer, equipped with a TCI cryoprobe) on samples dissolved in D^2^O, to gain more information on the aliphatic and aromatic resonance regions. Backbone φ and ψ torsion angle constraints were obtained from a database search procedure on the basis of backbone (^15^N, HN, Hα) chemical shifts using TALOS+ (32). Hydrogen bond restraints were derived using standard criteria on the basis of the amide ^1^H/^2^H exchange experiments and NOE data. When identified, the hydrogen bond was enforced using the following restraints: ranges of 1.8–2.0 Å for d(N-H,O), and 2.7–3.0 Å for d(N,O).

A total of 600 three-dimensional structures were generated for I27 single-module from 1188 NOEs, 23 hydrogen bonds and 140 angular restraints using the torsion angle dynamics protocol of CYANA 2.1. The 20 best structures (based on the final target penalty function values) were minimized with CNS 1.2 according the RECOORD procedure (33) and analyzed with PROCHECK (34). Root Mean Square deviations were calculated with PYMOL (35). All statistics are given in Table 1 of Supplemental Material.

For I27 tandem, because the NMR resonances for residues at similar position in the N- and C-terminal domain were found strictly identical (see Results section), the same restraint set was used for the modeling of each module. Since this restraint set was found virtually identical to the one collected for I27 single-module, the 3D structure of each module in the tandem was found virtually identical to that of the single-module. Moreover, no additional long range nOe’s were found neither between the two modules nor between residues belonging to the segment tethering them, precluding any defined spatial relative positioning of the two modules.

### High-Pressure NMR Spectroscopy

To permit unfolding of the protein within our accessible pressure range (1-2500 bar), 1.7M guanidinium chloride was added to the NMR samples (final volume: 350 µL). Reliable assignment of the amide cross-peaks in the HSQC spectra at the denaturant concentration used in the high-pressure experiments was ensured by guanidinium chloride titration. NMR spectra were acquired at 600 MHz using a 5 mm o.d. ceramic tube from Daedelus Innovations (Aston, PA, USA). Hydrostatic pressure was applied to the sample directly within the magnet using the Xtreme Syringe Pump also from Deadelus Innovations. Spectra analysis was performed using Gifa (30).

#### Steady-State Study

Series of 2D [^1^H-^15^N] HSQC spectra were recorded in steps of 200 bar, with a 2 hours relaxation time after every pressure change, to allow the protein to reach full equilibrium. Relaxation times for the folding/unfolding reaction were previously obtained from series of 1D NMR experiments recorded after 200 bar P-Jump, following the increase of the resonance band corresponding to the methyl groups in the unfolded state of the protein. The cross-peak intensities for the folded species were measured at each pressure, then fitted with a two-state model:

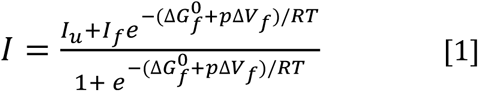

where I is the cross-peak intensity measured at a given pressure, I_f_ and I_u_ corresponds to the cross-peak intensity in the folded state (low pressure) and in the unfolded state (high pressure), respectively. ∆V_f_ and 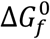 stands for the residue specific apparent volume change at equilibrium and free energy at atmospheric pressure, respectively. I_f_ and I_u_ values were floating parameters in the fit, and the data and fitted values were normalized (after the fit) using these plateau values to yield plots of fraction folded as a function of pressure.

Native contact maps were obtained by using software CMView (36) with a threshold of 9 Å around the Cα of each residue, using the best structure among the 20 refined ones. Using the geometric mean, rather than the joint probability as previously done (22,26), ensures the correct unfolding profile in the case of two-state unfolding.

#### Kinetics Study

A time series of [^1^H-^15^N]-SOFAST HMQC (37–39) were recorded for 2 hours with acquisition time of 2 min (4 scans with a relaxation delay of 125 ms; time domain in the indirect dimension: 64 complex points) after each positive 200 bar pressure jump. The jump took about 15 s, such that no dead time was apparent in the relaxation profile. The pressure-jump relaxation profiles at each pressure were well fitted with a single-exponential decay model.

For a simple two-state reaction, the P-jump relaxation time τ is the inverse sum of the folding and unfolding rates (τ_(p)_ = 1/(k_u(p)_ + k_f(p)_)). The folding and unfolding rate constants (k_f_ and k_u_) are exponentially dependent on the pressure through the activation volume for folding reaction (∆V‡_f_) and unfolding reaction (∆V‡_u_), respectively:

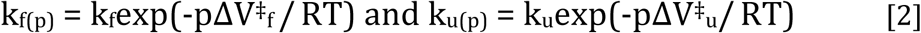

The pressure dependence of the ln(τ_(p)_) was fitted using a non linear least-squares analysis to extract the values of the activation volume and folding/unfolding rates at atmospheric pressure, using the following equation for τ_(p)_:

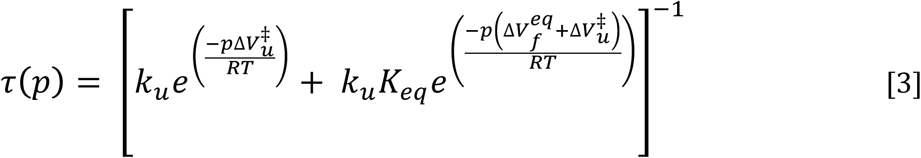

We constrained the ln(τ_(p)_) versus pressure fit using the previously determined equilibrium volume change 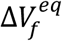 and free energy for folding 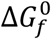 (see above) measured

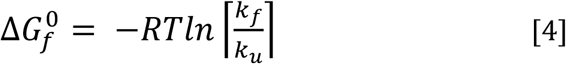

with Keq = k_f_/k_u_ and 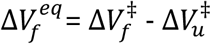.

### Proton/Deuteron exchange measurements

H/D exchange experiments were performed as previously described (40) with a 1 mM sample of I27 single-domain freshly dissolved in D_2_O. A series of [^1^H-^15^N]-HSQC spectra were recorded at either 600 MHz with a common dead time of 15 min and a time limit of 96 h. Amide proton protection factors (41) were calculated from the observed exchange rates (kex) obtained from the time dependence of the peaks intensities using an exponential decay.

## RESULTS

### NMR Resonances Assignment and Solution Structure of Titin I27

Proton and nitrogen NMR resonances of Titin I27 single-domain, renumbered to 1-91 for simplicity, have been assigned and its solution structure solved using essentially [^1^H,^15^N] double-resonance 3D NMR spectroscopy (see Material & Methods). Indeed, in the present study we used the construct previously described by Carrion-Vazquez et al. (15), also used by Rico et al. (18) that presents 3 mutations with regard to the protein which NMR structure has been published earlier (PDB code 1TIU) (42): V11A, T42A and A78T, as well as two additional residues at the C-terminal end (R90 and S91), used for the junction between the two modules in the tandem construct.

In addition, the structure of the I27 single-domain published by Improta et al. (42) was solved at acidic pH (pH 4.5, acetate buffer) and relatively high-temperature (35°C), although the present studies were conducted at neutral pH (pH 7.0, Tris buffer) and around ambient (25°C) temperature. This brings substantial chemical-shift perturbations in the 2D [^1^H,^15^N] HSQC spectrum, thus justifying its re-assignment (Fig 1). Once assigned, the ^15^N-edited 3D NOESY and TOCSY spectra, as well as a 2D NOESY spectra recorded at 800 MHz in a sample dissolved in D2O buffer, were used by the CYANA program to iteratively build models of the the solution structure. A final set of about 1300 restraints was used, and the pairwise RMS deviation calculated for backbone heavy atoms among the resulting 20 best refined structures was 1.08 ± 0.24 Å (Fig. 2, see Supplemental Material, Table 1 for the complete statistics).

**Figure 1.**
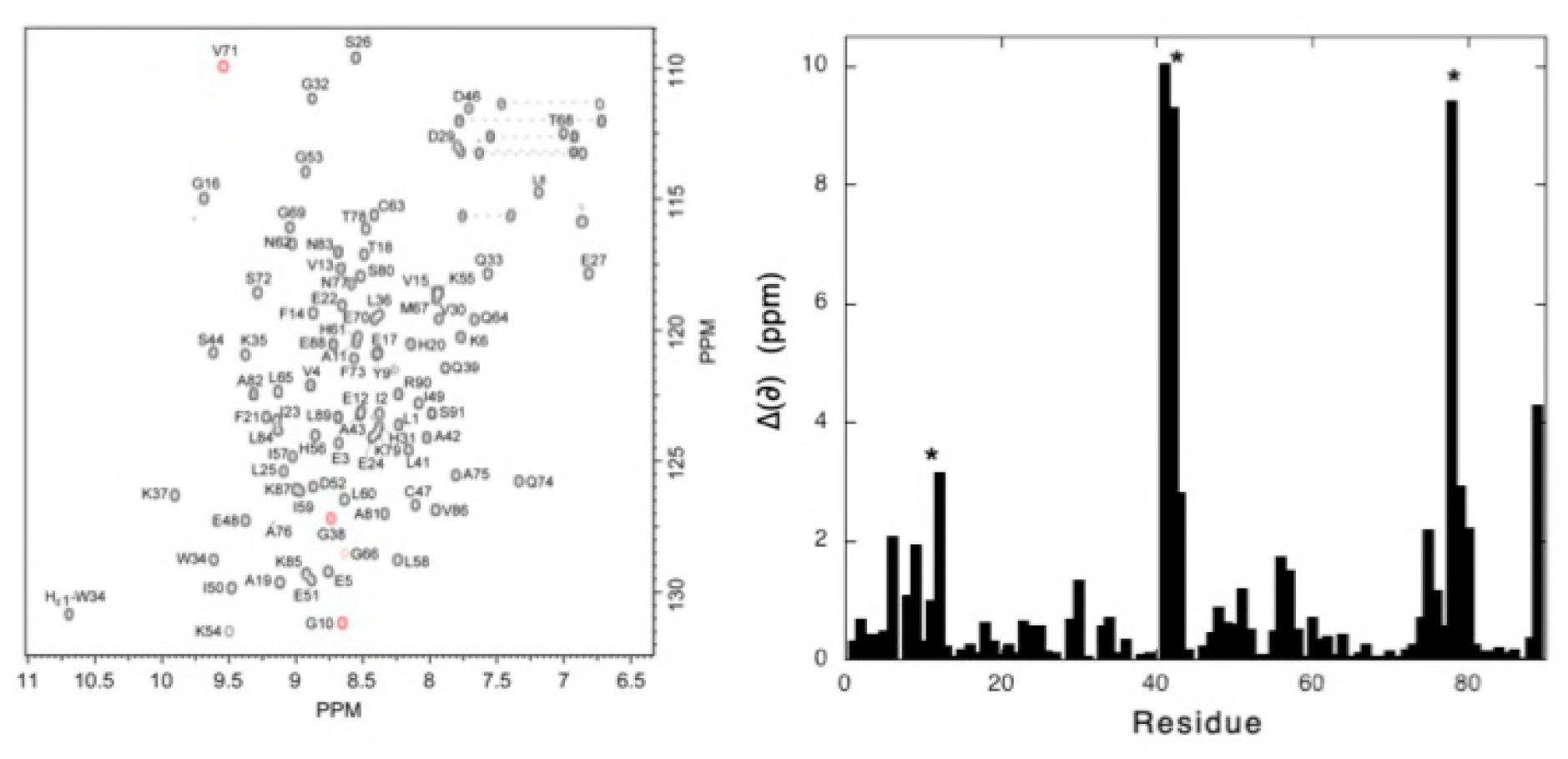
Titin I27 single-module NMR fingerprint. (Left) [^1^H-^15^N] HSQC spectrum of I27 single-module at 600 MHz, 25°C and pH 7.0 on a 1 mM, ^15^N-uniformly labeled sample dissolved in a 20 mM phosphate buffer (10 mM DTT). Cross-peak assignments are indicated using the one-letter amino acid and number. Cross-peaks in red are for amide groups with aliased ^15^N resonances. (Right) Average amide chemical-shift variations, calculated as 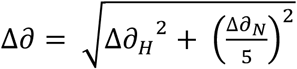, measured between the I27 single-module construct used in this study and the variant which NMR structure has been previously published by Improta et al. (1996). The stars indicate the mutations between the two constructs.

**Figure 2.**
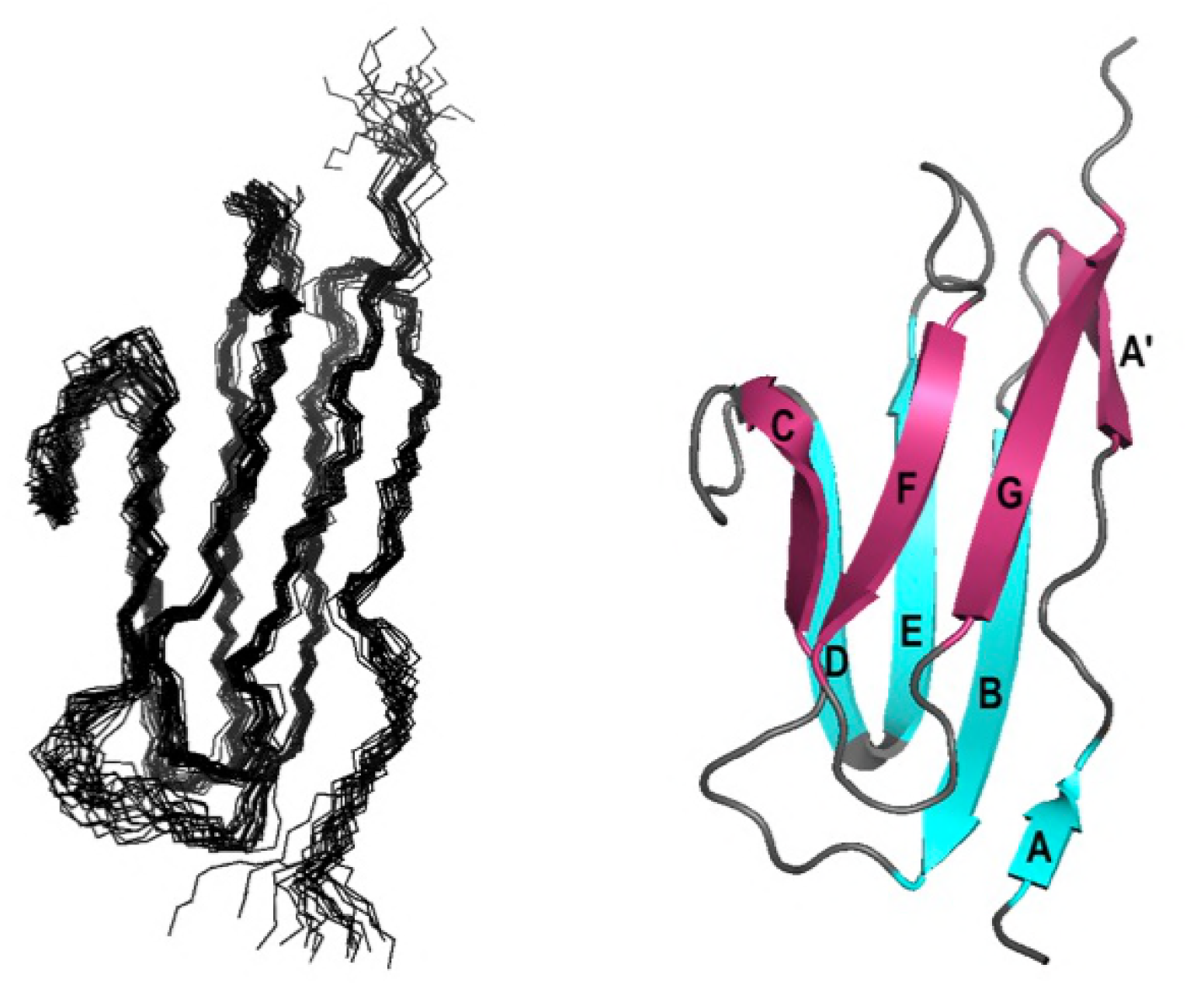
Solution structure of titin I27 single-module. (Left) Overlay of the 20 best NMR structures with lowest energy. (Right) Ribbon representation of the solution structure of I27 single-module. The two ß-sheets forming the ß-sandwich have been colored in blue and pink, and the different strands labeled using the standard nomenclature used for Ig-like modules.

As expected, we obtained the well-known Ig-like fold of the titin I-band Ig domains, that closely resembles to the previously published NMR structure (RMSD= 1.6 Å) or X-Ray structure (43) of a similar construct (PDB code 1WAA; RMSD=1.5 Å). It consists of a sandwich of two ß-sheets packing against each other (Fig. 2). Each of the sheets contains four strands: the first sheet comprises strands A, B, E and D, whereas the second sheet comprises strands A’, G, F and C. All adjacent strands are antiparallel, with the exception of the A’-G pair. Note that, the short strand A is poorly defined, due to strong distortions brought by the bulge involving residues Glu5-Lys6 and the proline at position 7. In the crystal structure, the AB sheet is stabilized by 3 H-bonds: E24(O)-K6(H), K6(O)-E24(H) and E3(O)-S26H. In our solution structure, only the amide protons of Lys6 and E24 appears to be (weakly) protected from solvent exchange (see further), suggesting that the third H-bond is not present or only transient (see further). In the following, the best calculated structure has been chosen to display the data gathered from the unfolding study.

Titin I27 tandem, renumbered for simplicity to 1-91 (first module) and 101 to 191 (second module) was also assigned. Surprisingly, in the 2D [^1^H,^15^N] HSQC recorded for I27 tandem, we observed a quasi-perfect overlapping with the HSQC recorded for the single-domain construct for all amide cross-peaks that corresponding to equivalent residues in the two concatenated modules (Supplemantal Material, Fig S1), with a slight line broadening, as expected from the increase of the molecular weight. Additional cross-peaks were observed only for residues L89, R90 and S91 in the first module, and residues L101, I102 and E103 in the second module. They correspond to the N- and C-terminal ends of the domain, free in the single-domain construct but linked in the tandem, hence the variations in chemical shifts. This suggests that the two modules behave independently in the tandem, without any strong interactions between the two modules. This was further confirmed by the inspection of the NOESY spectra, where no interaction can be detected between the two modules.

In the following, in the case of I27 tandem we will consider only the residues that give rise to overlapping cross-peaks in the HSQC spectra of the single-domain and of the tandem repeat. Indeed, no long-range nOe’s have been detected within the peptide segment L89-R90-S91-L101-I102-E103 linking the two modules, indicating that it does not adopt any defined conformation. Since this segment is not structured, it is unlikely to contribute to ∆V.

### High-Pressure Unfolding Monitored with NMR Spectroscopy: Steady-State Study

2D [^1^H,^15^N] HSQC spectra of ^15^N uniformly labeled titin I27 single-module and tandem in the presence of 1.7 M guanidinium chloride were recorded as a function of pressure (Fig. 3). Without denaturant, we cannot unfold the protein in the pressure range allowed by our experimental setting: this was expected since the I27 domain was found already to be extremely stable against temperature and chemical denaturation (13). In that case, we observed only pressure dependent shifts of the native-state resonance frequencies, due to compression and possible population of higher energy low-lying states (44–45). Interestingly, in the case of the tandem, we do not observe any separation for the cross-peak corresponding to equivalent residues in the two modules: whatever the pressure where the measurement was done, the HSQC spectra of the single-module and of the tandem overlap. This was also true when adding the denaturant, but, in addition, we observed a decrease in overall intensity of each native state peak as a function of pressure. Concomitantly we observed an increase in the intensity of the peaks centered around 8.5 ppm in the proton dimension, due to population of the unfolded state. These observations indicate that each residue of titin is in slow exchange between the chemical environments it experiences in the folded and unfolded states. Hence, the loss of intensity of the native state resonances reflects directly the decrease in population of the folded state as detected locally by each residue.

**Figure 3.**
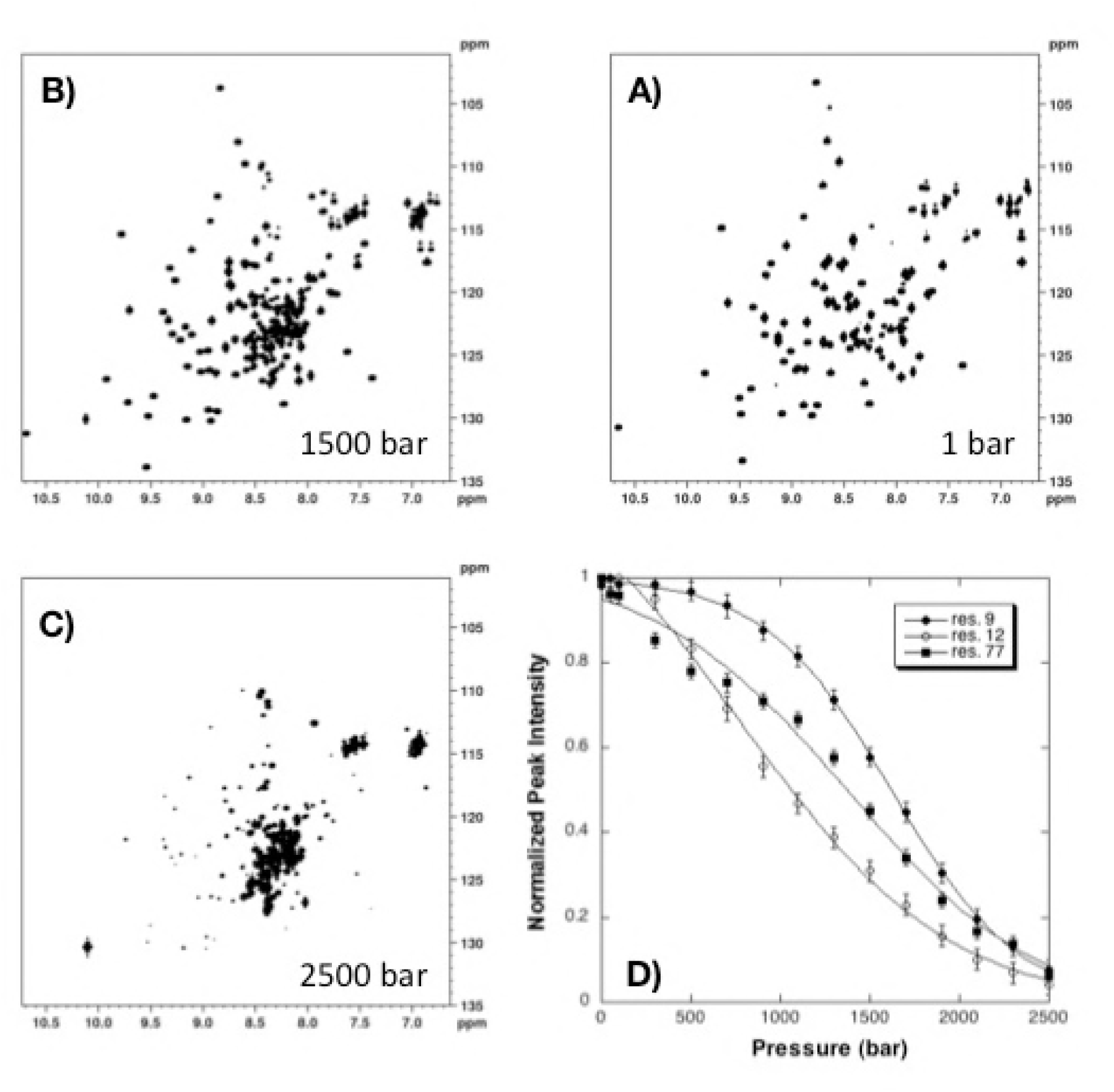
NMR detected high pressure unfolding of I27 single-module at 298 K and 1.7 M guanidinium chloride. (A-C) Examples of [^1^H-^15^N] HSQC NMR spectra at different pressures as indicated; (D) Examples of 3 residues exhibiting distinct unfolding profiles. A similar behavior was observed for I27 tandem (not shown).

This allowed us to fit the local pressure unfolding curves, obtained from the decrease in intensity of each individual peak for all resolved resonances individually to a two-state pressure-induced unfolding model as described in Materials and Methods (Equ [1], Fig 3), yielding residue specific values for the apparent volume change (∆V_u_ = -∆V_f_) and apparent free energy (∆G_u_) of unfolding (22) (Fig. 4).

**Figure 4.**
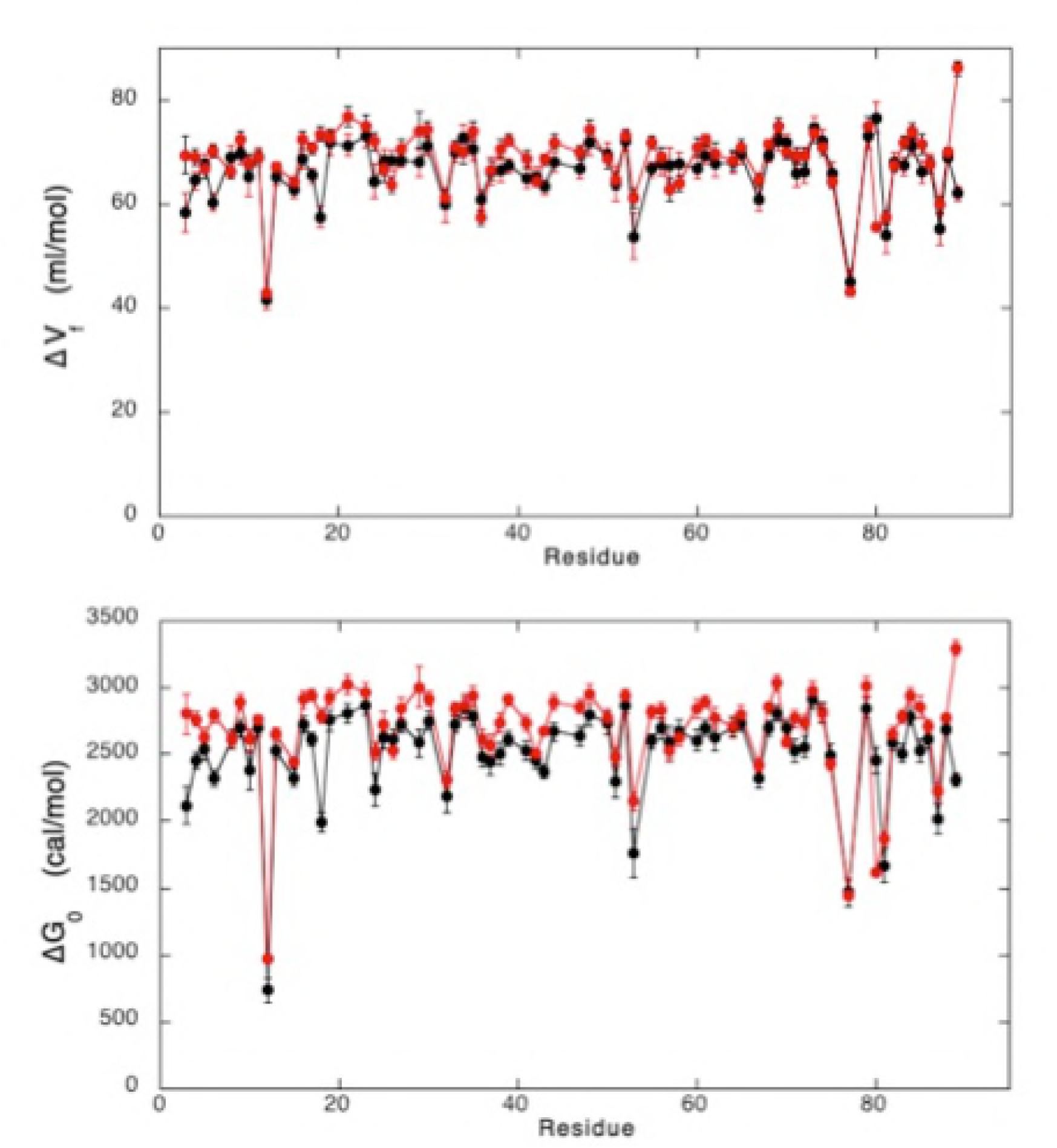
Steady-state thermodynamic parameters. Apparent residue-specific ∆V_f_ (top) and ∆G_0_ (bottom) values obtained through the fit of the intensity decrease of the 2D [^1^H-^15^N] HSQC cross-peaks with pressure recorded on I27 single-module (black dots) and tandem (red dots), plotted versus the protein sequence. For I27 tandem, the data corresponds both to residue 1-91 and 101-191, Since residues at similar position in the N- and C-terminal modules in I27 tandem give rise to unique amide cross-peaks, they also share the same ∆V_f_ and ∆G_0_ values. For simplicity, the same x-axis (residues 1 to 91) has been used for the two constructs.

At 1.7 M guanidinium chloride, titin I27 exhibits moderate stability, with average values <∆G_u_> = 2.45 ± 0.17 kcal/mol for the single module and 2.69 ± 0.06 kcal/mol for the tandem. Similarly, average <∆V_u_> values of −66.19 ± 2.21 ml/mol and −68.7 ± 1.55 ml/mol were obtained for the two constructs, respectively. Note that whether the two-state model was adequate for all individual unfolding profiles, distinct unfolding profiles for different residues were observed, demonstrating clear deviation from two-state behavior of the global unfolding for the two constructs. This leads to asymmetric distributions of apparent ∆V_f_ (Supplemental Material, Fig S2) and apparent ∆G_f_, suggesting that regions in the I27 domain may unfold at lower pressures.

To visualize more precisely which regions of the protein become disordered at intermediate pressures, we constructed fractional contact maps (22) for I27 single-module. We define the probability of contact for any pair of residues, P_i,j_, as the geometric mean of the fractional probability that each of the two residues is in the folded state at a given pressure (P_ij_ = [P_i_ x P_j_]^1/2^) (26). The pressure dependence of the contact maps showed that the region first affected by an increase of pressure is the short parallel ß-sheet (Strands A’G). A significant (P_ij_ ≤ 50%) loss of contacts in this ß-sheet is apparent already at 600 bar (Fig. 5). Between 600 and 1000 bar, it extends to neighboring residues in the C-terminal pole of the ß-sandwhich. At 1200 bar, it concerns most of the residues in the I27 domain, as a result of water penetration in the whole hydrophobic core of the molecule and disruption of the 3D structure. Because similar values and similar distribution of ∆V_u_ were measured for I27 modules in the tandem construct, similar results can be drawn.

**Figure 5.**
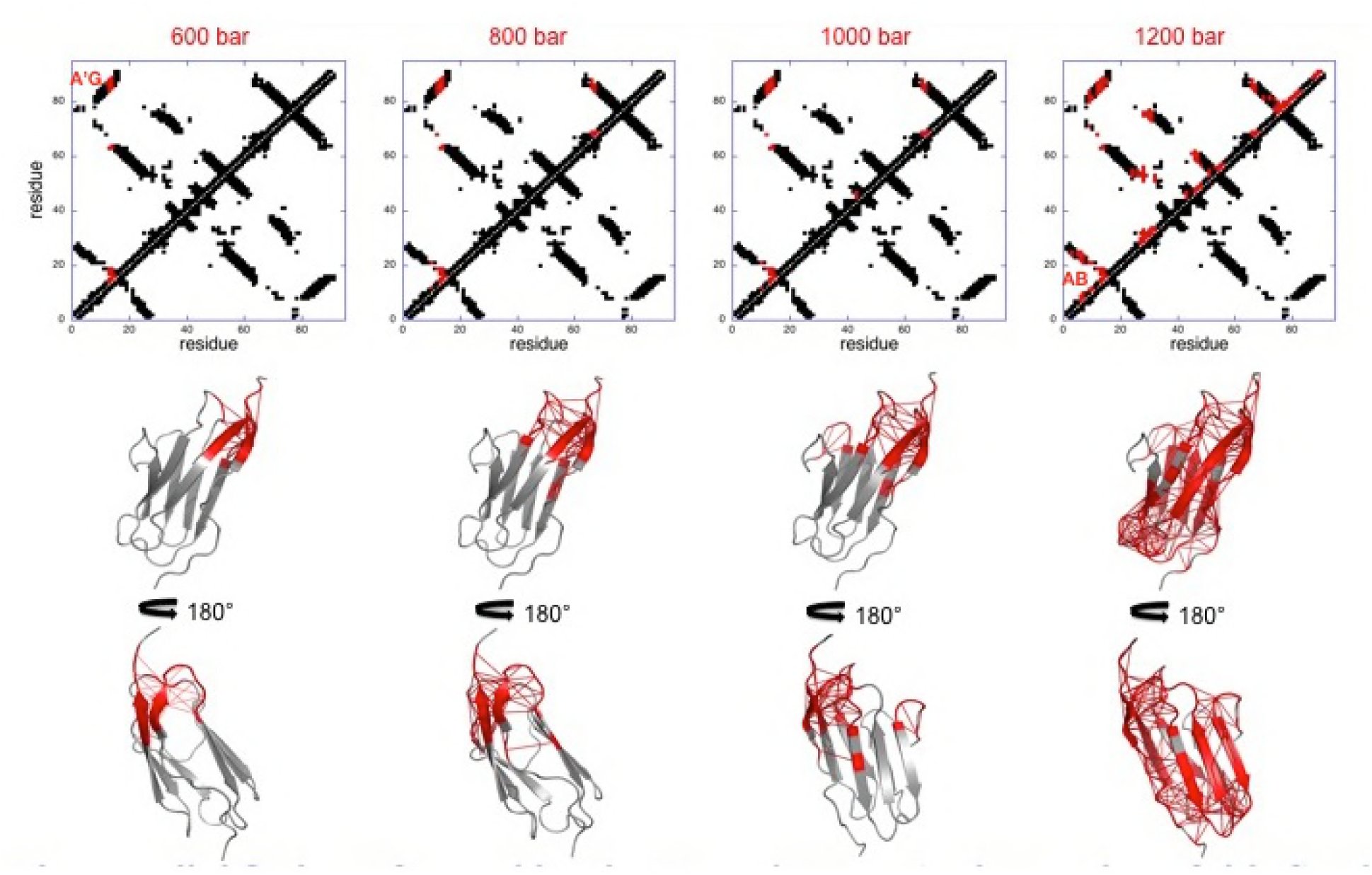
Pressure denaturation of Titin I27 single-domain. (Top) Contact maps built from the best solution structure obtained for I27 single-module at 600, 800 1000 and 1200 bar, as indicated. The contacts above the diagonal have been colored in red when contact probabilities (P_ij_) lower than 0.5 is observed. (Bottom) Ribbon representations of the solution structure of I27 where the red thicks represent contacts that are weakened (P_ij_ ≤ 0.5) at the corresponding pressures.

### High-Pressure Unfolding Monitored with NMR Spectroscopy: Kinetics Study

Real-time pressure-jump NMR Spectroscopy was used to probe further the folding reaction of both titin I27single-module and tandem, as previously described (24). A total of 60 2D [^1^H,^15^N] SOFAST-HMQC, each of 2 min acquisition time, were collected as a function of time after positive pressure jumps of 200 bar. Despite the weak decrease in intensity during the relaxation time (Fig 6), yielding relatively large experimental errors, the time dependence of each cross-peak intensity was well described by a single-exponential time (τ). The dependence of ln(τ) versus pressure chevron plots obtained for each residues were analyzed to yield residue-specific activation volume of folding (∆V_f_‡) (Fig 6) that give access to the hydration state of the specific residue at the Transition State Ensemble (TSE).

**Figure 6.**
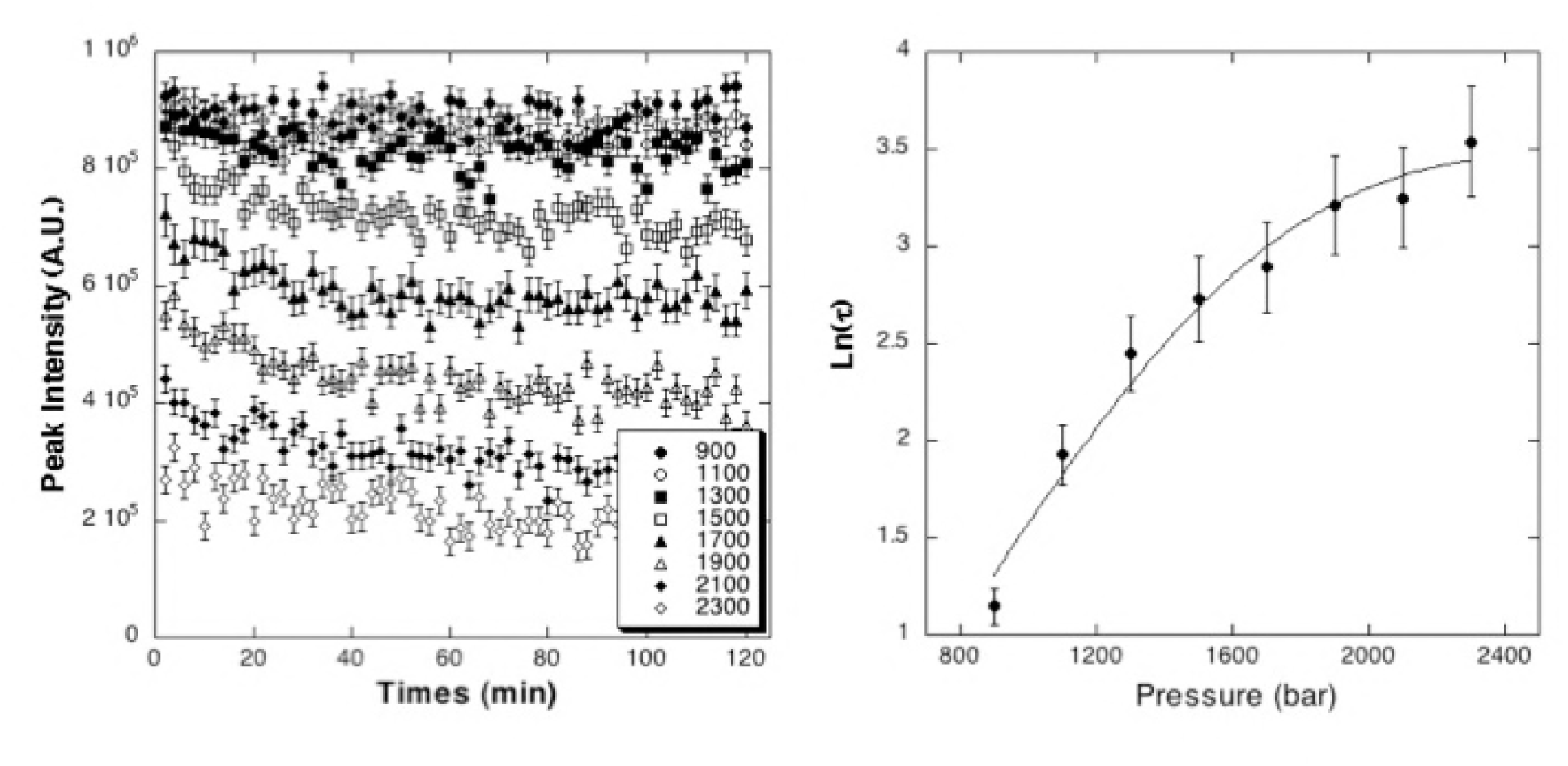
P-Jump analysis on Titin I27 single-domain. (Left) Time evolution of the amide cross-peak intensity for a representative residue (K79) after 200 bar positive P-jumps realized at different pressures (final pressures reported in the insert). The curves were fitted with exponential decays to obtain the values of τ. (Right) Chevron plot of ln(τ) versus pressure. The fit with Eq [3] allows extracting the folding and unfolding rate constants k_u_ and k_f_, as well as the apparent residue-specific activation volume of folding (∆V_f_‡) (See Material and Methods).

The analysis of the distribution of the hydrated/dehydrated residues within the TSE gives in turn a good picture of the distribution of the cavities formed at the TSE. Unfortunately, the uncertainties on τ values were too large to assess accurately any structural heterogeneity in the cavity distribution at the TSE, as it has been done previously for others proteins (24). Nevertheless, from ∆V_f_‡ mean values calculated over all the residues, we were able to demonstrate that the volume of the TSE is very close to the folded state for both constructs (Fig 7). Mean values of ∆V_f_‡ = 65.77 ± 6.33 ml/mol and 67.73 ± 7.52 ml/mol were measured for titin I27 single-module and tandem, respectively, very close to those measured for ∆V at equilibrium. This means that most of the cavities present in the folded state are already formed in the TSE in both constructs, in other words that the TSE is very dehydrated. Average kinetics constant of unfolding at ambient pressure can also be obtained from the fit: k_u_^0^ = 6.6 (± 3) x 10^−4^ s^-1^ and 4.6 (± 1.6) x 10^−4^ s^-1^ can be estimated for I27 single-module and tandem, respectively, in good agreement with rate constant values previously obtained for similar titin Ig single-module or tandem repeat (16, 46).

**Figure 7.**
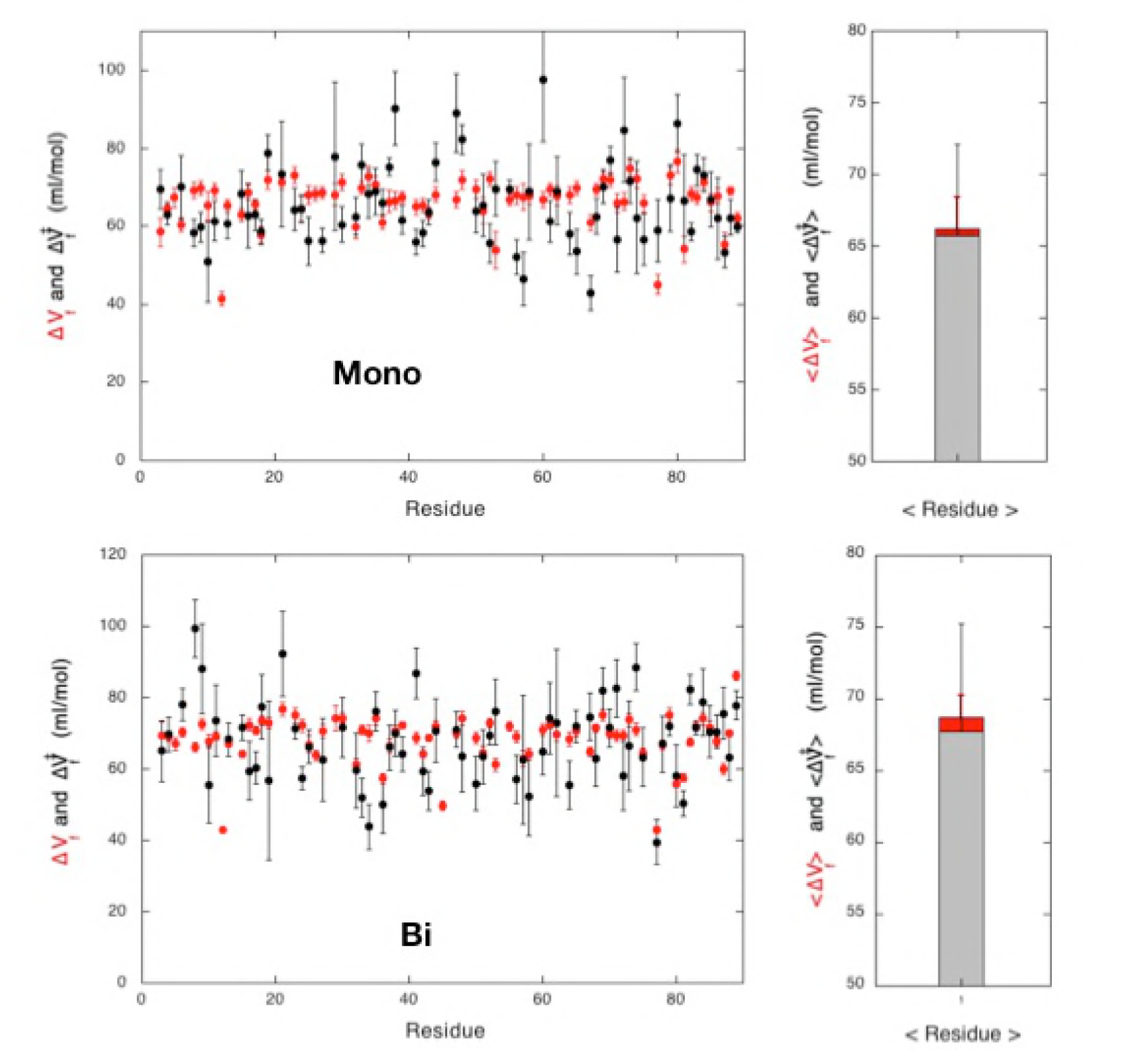
Kinetics Parameters of unfolding. (Left) Apparent residue-specific ∆V_f_ values measured at equilibrium (red dots) and ∆V_f_‡ values obtained from P-Jump experiments (black dots) for titin I27 single module (Top) and tandem (Bottom) are plotted versus the sequence. As in Figure 4, the same x-axis (residues 1 to 91) has been used for the two constructs, for simplicity. (Right) Average values of ∆V_f_‡ (dashed bar) and ∆V_f_ (red bar) calculated over all the residues for I27 single-module (Top) and tandem (Bottom).

### Local Stability Probed by Proton/Deuteron Exchange

H/D exchange for the single I27 domain of titin was investigated in our experimental conditions (pH 7 and T=298K), but in the absence of chemical denaturant, to explore the local stability of the domain. To do this, [^1^H,^15^N] HSQC were recorded with time on a protein sample freshly dissolved in D_2_O (See Materials and Methods for details). Accurate fit with an exponential decay can be obtained for the HSQC cross-peak intensity of 41 residues over 88 non-proline residues, and the corresponding protecting factors (PF) were calculated (41) (Supplemental Material, Fig S3). The local stability of the different ß sheets was then estimated by the average value of the protecting factors (<PF>) calculated over the amide protons involved in the H-bond stabilizing the ß-sheet. A very low <PF> value was measured for the AB antiparallel ß-sheet, probably due to a frequent opening of this ß-strand pair, as it was suggested already (42, 47).

With the exception of this highly distorted antiparallel AB ß-sheet, the less stable sheet appeared to be the short parallel ß-sheet formed by strands A’ and G, with an average PF value smaller than about one order of magnitude when compared to the values calculated for the antiparallel ß-sheets forming the rest of the ß-sandwich (Fig 8). This can be explained by the fact that H-bonds in parallel ß-sheets are known to be less stable than in antiparallel ß-sheets, due to a less regular alignment of their constitutive atoms (C-O---HN), hence more prone to solvent exchange.

**Figure 8.**
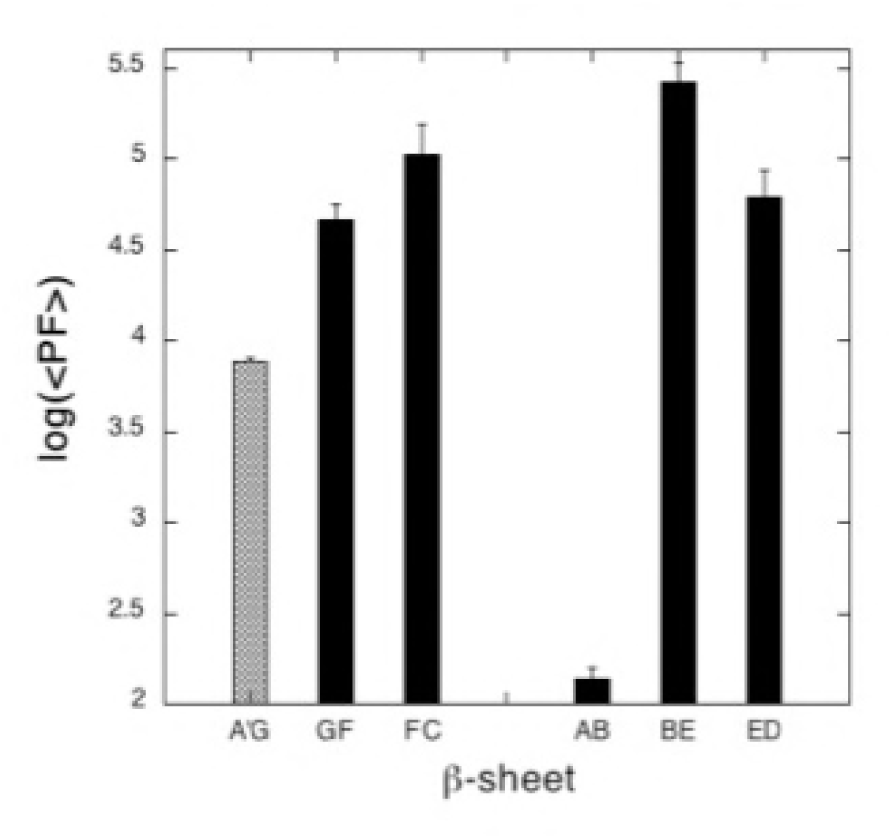
Average values <PF> calculated for amide protection factors stabilizing each double-stranded ß-sheet in the ß-sandwich of titin I27 domain. The dashed bar stands for the A’G parallel ß-sheet, the dark bars for antiparallel ß-sheets.

## DISCUSSION

In this study we have monitored the pressure unfolding of titin I27 single-module and tandem with high-resolution NMR spectroscopy. We used the same constructs already studied with force spectroscopy, with the advantage of being able to compare directly the results of these different approaches. Nevertheless, this choice implies to reassign the NMR spectra of our constructs and to check the integrity of their solution structures in our experimental conditions. Indeed, assignment and solution structure were available only for a construct of I27 presenting 3 mutations when compared to our single-module construct, and at higher temperature and acidic pH (42). As expected, we found that I27 keeps it Ig-like fold at pH 7 and 25°C, even though the short AB ß-sheet appears loosely defined, due to the presence of a ß-bulge already reported in the previous NMR and X-rays structures (43).

Interestingly, the local unfolding of this short ß-sheet has been proposed as the starting event of titin Ig domains unfolding, yielding an intermediate folding state where strand A is detached from the ß-sandwhich. This folding intermediate was first observed in modeling studies (48), confirmed by careful analysis of AFM traces (14), and has been characterized by biophysical and structural techniques (49). Thus, steered molecular dynamics simulations (SMD) (50–53) on titin I27 domain revealed that the source of titin’s secondary structure elasticity lays in the architecture of its N-terminal ß strands (48; 54-58). These simulations showed that I27’s unfolding comes about through nearly synchronous breaking of a network of hydrogen bonds that stabilize the terminal ß strands of the domain. They also showed that the rupture of the AB ß-sheet was a minor event, without major contribution to mechanical stability, preceding the rupture of the A’G patch yielding the full unraveling of the module. These simulations are in good agreement with the ≈ 6 Å elongation observed with the AFM at forces ≈ 100 pN, that corresponds to the stiffening of the N-terminal strand and neighboring residues after breaking of the short AB ß-strand, while rupture of the A’G sheet is visible at higher forces in AFM experiment (≈ 200 pN). This was also corroborated by force spectroscopy studies performed on mutants where interactions between AB strands were weakened (14). Thus, the two small ß-sheets AB and A’G guard the titin domain against forced unfolding like two pieces of “molecular Velcro”. But the A’G sheet appeared to be crucial for mechanical resistance, forming a mechanical clamp (14, 59).

However, the intermediate folding state is generally not observed in chemical denaturation studies (10), and was not detected also in the present high-pressure NMR study: the first event in I27 unfolding was found to be the rupture of the A’G parallel strand. The local pressure disruption of this sheet starts around 600 bar and concerns mainly this secondary structure at pressures up to 1000 bar. Beyond 1000 bar, all the protein unfolds, including the AB sheet. This is due to water penetration into the hydrophobic core of the molecule, probably through the breach created by the disruption of the A’G ß-sheet. Before to conclude in a striking difference between the effect of high hydrostatic pressure and stretching, it should be mentioned that the existence of this intermediate remains controversial: from AFM results obtained after single conservative amino-acid mutations of the I27 domain, Williams et al. (59) have postulated that this intermediate is not populated under physiologically relevant conditions (forces <100 pN), and, in a recent study, using ultra-stable magnetic tweezers Chen et al. (46) cannot identify this intermediate state as well.

Also, this difference of behavior observed between the effect of high hydrostatic pressure and stretching might originate from the difference in the forces at play. When using force spectroscopy, the molecule is stretched while maintained at its N- and C-terminal ends. By the way, it is conceivable that the N-terminal segments breaks sequentially, starting from the less stable secondary structure (the AB sheet) to the more stable one (the A’G sheet). From H/D exchange experiments, we showed that the AB sheet is indeed the less stable structure, with the weakest <PF> value. The A’G parallel sheet appears more stable, even if the <PF> value measured for this sheet is well below those measured for the others antiparallel ß-sheets forming the ß sandwich. High hydrostatic pressure unfolds protein by pushing water molecules inside void cavities in the core of the molecule. Due to their weaker stability, the AB and AG’ sheets provide attractive entry points for water. Why choosing the AG’ sheet first instead of the AB sheet of weaker stability? This point remains unclear even though it seems corroborated by recent results reported by Stacklies et al. (43). Using MD simulations to analyze the force distribution inside I27 after applying force, they showed that the force was directly deflected into the protein core via mainly hydrophobic side chain interactions between strands A and G, bypassing the AB inter-strand hydrogen bonds. Moreover, this observation is also supported by a previous ϕ value analysis performed by Fowler and Clarke (10) on titin I27 domain. They showed that only the A’ and G strands were completely unstructured in the transition state of I27, with ϕ values lower than 0.2, while all other strands of the ß sandwich are structured to some extent.

Our pressure-jump experiments also clearly demonstrate that the transition state ensemble (TSE) of titin I27 single-module is highly dehydrated, with an average activation volume ∆V_f_‡ found very close to the ∆V value measured at equilibrium. That means that most of the cavities present in the folded state are already formed in the TSE. Nevertheless, the measurement of the relaxation time values (τ) were not accurate enough to determine the precise location of these cavities in the TSE, precluding to assess if the A’G is hydrated in the TSE, as suggested by the ϕ value analysis. Besides, residue-specific unfolding constant rate can be extracted from the chevron plots that were found in good agreement with previous studies (16, 18, 46).

When regarding titin I27 tandem, our results clearly confirm previous findings: I27 tandem behaves “as the sum of its parts” (16). No interaction can be found between the two modules whose structure in solution remains identical to that described for I27 single-module. Indeed, identical chemical shifts were measured for residues at similar position in the two tethered modules, yielding a quasi-perfect overlapping whatever the conditions of the study (concentration of guanidinium chloride, pressure). Moreover, only a slight line broadening was observed in the tandem, suggesting that the two modules move independently. A similar behavior was observed also for the two constructs upon unfolding, suggesting a similar folding landscape and similar folding pathways for titin I27 domain when tethered or untethered. As previously reported for others titin Ig domains (16), we observed nevertheless a very small increase in stability on extension of the isolated domain to the tandem. Similarly, unfolding kinetics of I27 single-domain appear slightly slower than in the tandem. Nevertheless, the differences measured in the present study are in the range of experimental errors, making it difficult to conclude about their relevance.

## CONCLUSION

In summary, without the help of MD or protein engineering, we were able to experimentally decipher the molecular events leading to the unfolding of titin I27 single- or tethered module. This is a result of the residue-specific analysis allowed by high-pressure multidimensional NMR: it gives local, residue-specific information of what happen to the protein structure during the protein unfolding process. This is very different from force spectroscopy approaches, for instance, that give only indirect clues on the unfolding process, like the magnitude of the forces at play or the global stiffening experienced by the structure under stretching. Such global information must be corroborated by MD simulations and/or by the evaluation of site-specific mutation effects in order to be converted into an atomic description of the mechanism of unfolding. By a direct and experimental measurement of unfolding representative data at a residue level (apparent ∆G, ∆V, ∆V_f_‡, folding and unfolding rates), high pressure NMR allows to observe the sequential events during unfolding at a molecular level.

## ACKNOWLEDGEMENTS

The authors gratefully acknowledge Frederic Allemand (CBS, Montpellier) for the sub-cloning of the constructs used in this study. All members of the CBS are supported by the French Infrastructure for Integrated Structural Biology (FRISBI) ANR-10-INBS-0.5.

